# The mammalian CLU homolog *FMT* controls development and behavior in *Arabidopsis*

**DOI:** 10.1101/2020.08.01.231183

**Authors:** Alexandra Ralevski, Federico Apelt, Justyna J. Olas, Bernd Mueller-Roeber, Elena I. Rugarli, Friedrich Kragler, Tamas L. Horvath

## Abstract

Mitochondria in animals are associated with development, as well as physiological and pathological behaviors. Several conserved mitochondrial genes exist between plants and higher eukaryotes. Yet, comparative mitochondrial function among plant and animal species is poorly understood. Here, we show that *FMT* (*<u>F</u>RIENDLY <u>MIT</u>OCHONDRIA*) from *Arabidopsis thaliana*, a highly conserved homolog of the mammalian *CLU* (*<u>CLU</u>STERED MITOCHONDRIA)* gene family encoding mitochondrial proteins associated with developmental alterations and adult physiological and pathological behaviors, affects whole plant morphology and development under salt stress and control conditions. *FMT* was found to regulate mitochondrial morphology and dynamics as well as germination, root length, and flowering time. Here, we show that it also affects leaf expansion growth, salt stress-responses and hyponasty. Strikingly, FMT impacted the speed of hyponasty with corresponding change in speed of locomotion of CLU heterozygous knockout mice. These observations indicate that homologous genes affect homologous functions in plants and animals offering the possibility to develop plant models for the study of mammalian behaviors.

## INTRODUCTION

Mitochondria function is critical for the health and lifespan of an organism, as they regulate vital cellular processes, including metabolism, energy production and apoptosis, and can alter their number, function or morphology in response to various environmental or cellular conditions (Chan, 2006; Mishra and Chan, 2016; Verdin et al., 2010). Mitochondrial dysfunction due to defects in changes in fission/fusion dynamics, altered trafficking, mutations of mtDNA, or impaired transcription, can lead to bioenergetic defects and an overall decline in the fitness of an organism. Dysfunctional mitochondria can induce alterations during development, as well as physiological and pathological malfunctions in adult organisms, for example in various neurodegenerative diseases, including Alzheimer’s disease (AD), Parkinson’s disease (PD), Huntington’s disease (HD) and schizophrenia (SCZ) (Bertholet et al., 2016; Burte et al., 2015; Golpich et al., 2017; Goncalves et al., 2015; Islam, 2017; Krols et al., 2016; Oka, 2016; Rajasekaran et al., 2015). Many proteins that regulate these critical mitochondrial processes are highly conserved between species, including plants and animals.

Changes in mitochondrial morphology and dynamics can elicit changes in cell functioning, which then can alter numerous physiological processes that produce changes at the level of the whole organism, such as phenotypic and behavioral changes (Bertholet et al., 2016; Burte et al., 2015; Golpich et al., 2017; Goncalves et al., 2015). Although this hierarchy of behavior is conserved among all species, plants are often excluded as exhibiting these conserved, fundamental behaviors. However, behavior simply derives from the response of an individual to its environment (Silvertown and Gordon, 1989). Thus, uncovering the cellular and molecular mechanisms that drive various behaviors in different organisms, especially those that are highly functionally conserved, is critical to unravel and examine fundamental principles that underlie physiological and pathological states across multiple species.

The plant model organism *Arabidopsis thaliana* and animals share numerous homologous genes encoding mitochondrial proteins that are functionally conserved, such as those that encode the fission proteins fission 1 (FIS1) and dynamin-related protein (DRP) (Aung and Hu, 2012; Scott et al., 2006; Zhang and Hu, 2009; Zhang and Hu, 2008), voltage-dependent anion channels (VDACs) (Robert et al., 2012), the mitochondrial calcium uniporter (MICU) (Wagner et al., 2015) and CLU, a member of the clustered mitochondria superfamily. Initially, CLU was characterized in slime mold (*Dictyostelium discoideum*) (Zhu et al., 1997) and yeast (*Saccharomyces cerevisiae*) (Fields et al., 1998), deriving its name from the striking phenotype of mitochondrial clustering following a gene knockout (KO), and was proposed to regulate mitochondrial orientation within the cell. However, recent evidence suggests a highly dynamic role for CLU in multiple mitochondrial processes. CLU in flies (*Drosophila melanogaster*) negatively regulates the interaction between PTEN-induced putative kinase 1 (PINK1) and Parkin necessary for mitochondrial autophagy (Sen et al., 2015). PINK1 and Parkin are mutated in early-onset Parkinson’s disease in humans (Hu and Wang, 2016). *CLU HOMOLOG* (*CLUH*) in mice regulates mitochondrial metabolism by specifically binding a subset of mRNAs encoding mitochondrial proteins (Gao et al., 2014, Schatton et al., 2017). Knockout of the *CLU* homolog in *Arabidopsis, FMT* (*<u>F</u>RIENDLY <u>MIT</u>OCHONDRIA*), produced alterations in mitochondrial trafficking and matrix exchange (El Zawily et al., 2014).

*CLU* is upregulated in the early stages of development in flies (Goh et al., 2013) and is upregulated in the liver at birth in mice (Schatton et al., 2017). *CLU* is also highly expressed throughout the body in mice with the highest expression in the gut, liver, kidney, heart and testis, and moderate expression in the forebrain and cerebellum (Schatton et al., 2017). However, the expression of *FMT*, a homolog of *CLU* in plants, remains poorly characterized. *CLU* is critical to regulate mitochondrial dynamics. Knockout of *CLU* in *Drosophila* has damaging effects– flies have a decreased lifespan of only 3-7 days, are sterile, and cannot fly (Cox and Spradling, 2009). Knockout of *CLU* in mice is lethal, as homozygous *clu*^-/-^ mice die shortly after birth and cannot be rescued (Schatton et al., 2017). Alternatively, while KO of *Arabidopsis FMT* causes phenotypic abnormalities (El Zawily et al., 2014), plants are still functional, mature to flowering, and produce seeds. Since *CLU* is highly conserved between plants and animals, *Arabidopsis* serves as an excellent model to study CLU function. Using this model system, we sought to elucidate the role of this gene, allowing for an understanding of its role in mitochondrial dynamics and behavior with implications for higher organisms.

## RESULTS

### Expression Pattern of *FMT* in *Arabidopsis*

We first investigated organ- and cell-specific expression of *FMT* in *Arabidopsis*. We performed RNA *in situ* hybridization on various tissue sections using an *FMT*-specific probe. *FMT* transcripts were present at a low level in the vegetative shoot apical meristem (SAM) and surrounding young leaves (Figure 1A), while *FMT* was highly expressed in the inflorescence and flower meristems (Figure 1B). *FMT* was also highly abundant in flowers, including carpels and stamens (Figure 1C-E). In addition, we found that *FMT* transcripts accumulated throughout the mid-stage embryo during the early and late torpedo stages (Figure 1G,H) but were absent from the mature embryo (Figure 1I). We included a control probe in the sense orientation, which gave no signal (Figure 1F). These data indicate a regulatory role for *FMT* in seeds and flower development in *Arabidopsis*.

**Figure 1:**
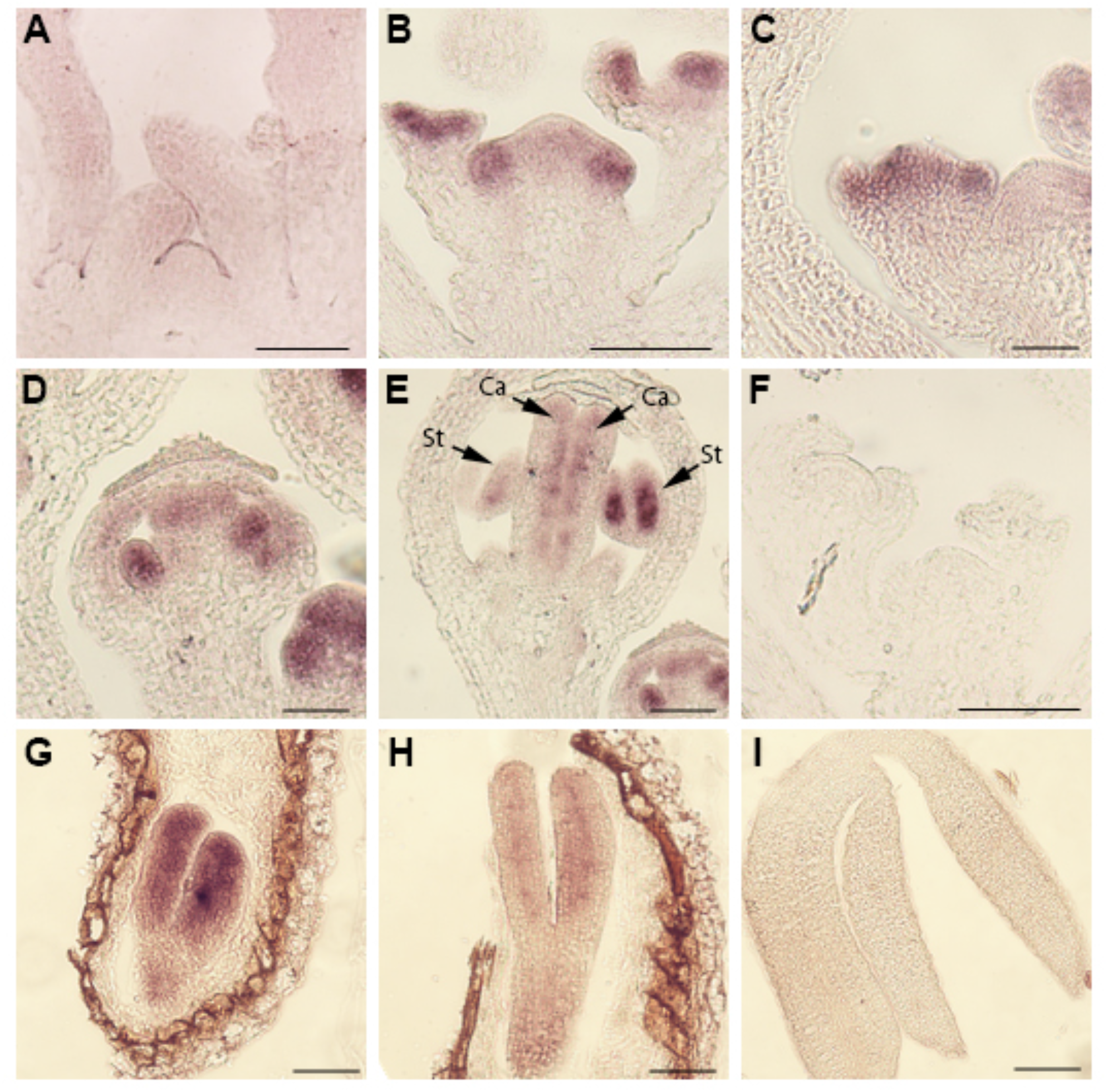
Expression pattern of *FRIENDLY MITOCHONDRIA (FMT)* in shoot apices, flowers and embryos of *Arabidopsis* Col-0 plants detected by RNA *in situ* hybridization. Longitudinal sections of vegetative meristem (A), inflorescence meristem (B), stage 4 flowers (C), stage 6 flowers (D) and stage 8 flowers (E), control probe in sense orientation in longitudinal sections of inflorescence meristem (F), embryos at various stages (G-I), early torpedo stage (G), late torpedo stage (H) and mature embryos (I). Arrow indicates *FMT* expression in stamen (St) and carpel (Ca) primordia. Scale bar = 100 μm.

### Generation of Overexpression Mutants of *FMT* in *Arabidopsis*

In order to assess the functional role of *FMT* in *Arabidopsis*, we created a transgenic plant line overexpressing (OE) *FMT* (referred to as *FMT*-*OE*) under the control of the cauliflower mosaic virus (CaMV) 35S promoter. We transformed *Arabidopsis* plants with *Agrobacterium* containing the pFASTG02-FMT overexpression vector. The pFAST vector carries a screenable GFP marker producing a signal detectable in the mature seed coat of transformed plants (Shimada et al., 2010). We selected transgenic seeds from transformed plants under a fluorescence microscope and identified homozygous T3 lines exhibiting 100% GFP fluorescence compared to wild-type (WT) seeds (Figure 2A-B). After creating a transgenic line overexpressing *FMT*, we obtained an *fmt* KO line from the Arabidopsis Biological Resource Center (ABRC, Ohio State University, Columbia, OH, USA). Genomic PCR verified this line contained an insertion at the first base of the second intron, preventing correct mRNA splicing (Logan et al., 2003) (Figure S1). qPCR determined the expression level of *FMT* in both the OE and KO lines. mRNA expression levels were almost two-fold higher in *FMT-OE* plants compared to the WT, while there was no detectable expression in the KO line (Figure S2).

**Figure 2:**
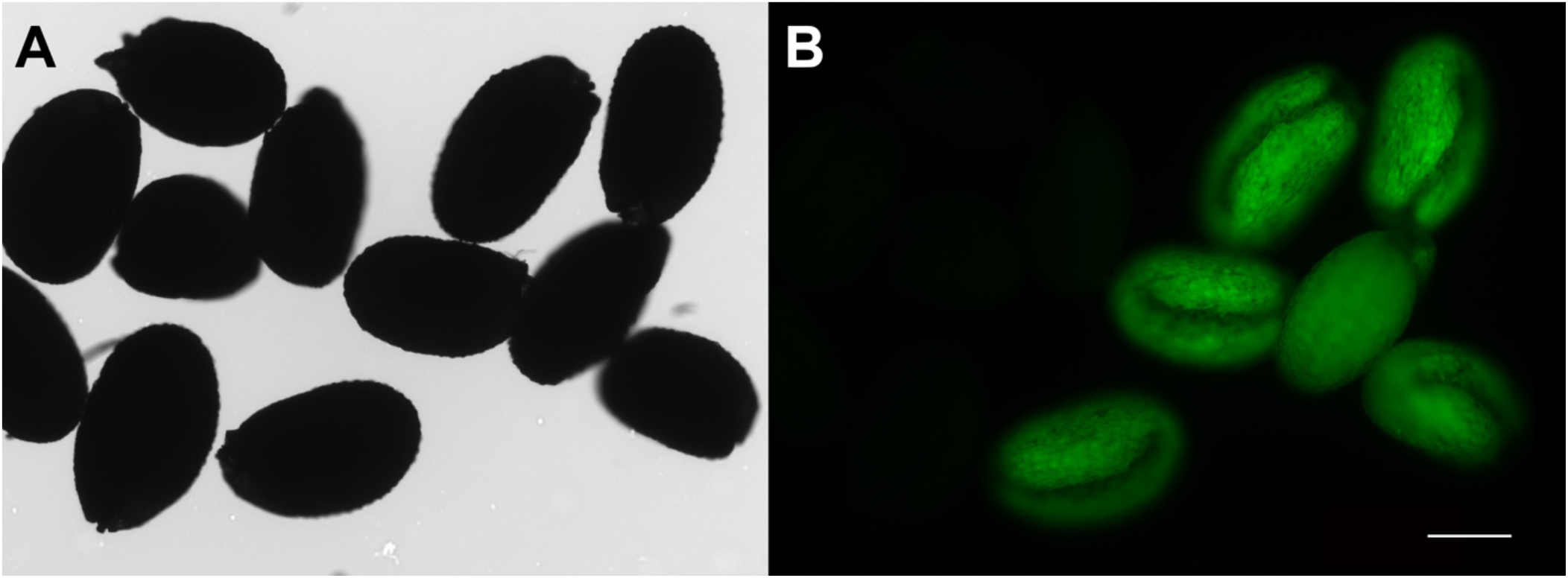
*Arabidopsis* seeds transformed with pFASTG02-FMT show green fluorescence. A) Transgenic seeds overexpressing *FMT* in the pFASTG02 vector show green fluorescence (right), while non-transformed WT seeds (left) do not show green fluorescence. B) Same field of view as in (A), viewed under bright field light. Scale bar = 100 μm.

### FMT Affects Mitochondrial Morphology and Dynamics

It was previously shown (El Zawily et al., 2014) that *fmt* mutants in *Arabidopsis* exhibit altered morphology, whereby organellar clustering is triggered by a lack of mitochondrial trafficking and motility, both known to be essential for proper cellular function. This phenotype of mitochondrial clustering often occurs under stress conditions, where mitochondria displayed altered motility and distribution (Chen and Chan, 2009; Nunnari and Suomalainen, 2012). Similar mutants with a non-functional *CLU* showed a significant increase in mitochondrial clustering in a variety of other species, including slime mold, yeast, fruit flies, and mice (Cox and Spradling, 2009; Fields et al., 1998; Gao et al., 2014; Schatton et al., 2017). Since *fmt* mutants have a significantly smaller root cap than WT plants (El Zawily et al., 2014), we analyzed mitochondria in root cap cells 7 days after germination (DAG) by transmission electron microscopy (TEM). We examined mitochondrial morphology and dynamics, including area, aspect ratio (AR), mitochondrial number, membrane fusions and clustering.

We found that plants lacking *FMT* showed a significant increase in mitochondrial number (Figure 3B,D), although AR and mitochondrial area remained unchanged (Figure 3E,F). WT and *FMT-OE* plants did not show a statistically significant difference in the number of mitochondria (Figure 3A,E,D). In order to analyze mitochondrial dynamics, we examined membrane interaction events of the inner mitochondrial membrane (IMM) and outer mitochondrial membrane (OMM) contacts in WT, *fmt* and *FMT-OE* plants. We found that *fmt* mutants displayed a significant increase in the number of IMM fusion events compared to both WT and *FMT-OE* (Figure 3G). In addition, *fmt* mutants also displayed a significant increase in the number of OMM fusion events compared to both WT and *FMT-OE* (Figure 3H). We then used the nearest neighbor value (Rn) to examine mitochondrial clustering in root cap cells. We found that mitochondria in plants lacking *FMT* trended towards clustering, while the mitochondria of WT and *FMT-OE* plants trended towards dispersion (Figure 3I). Taken together, our data confirms previous findings (El Zawily et al., 2014) and provides additional evidence that *FMT* regulates various mitochondrial mechanisms, including mitochondrial number, clustering, and inner and outer mitochondrial membrane fusions.

**Figure 3:**
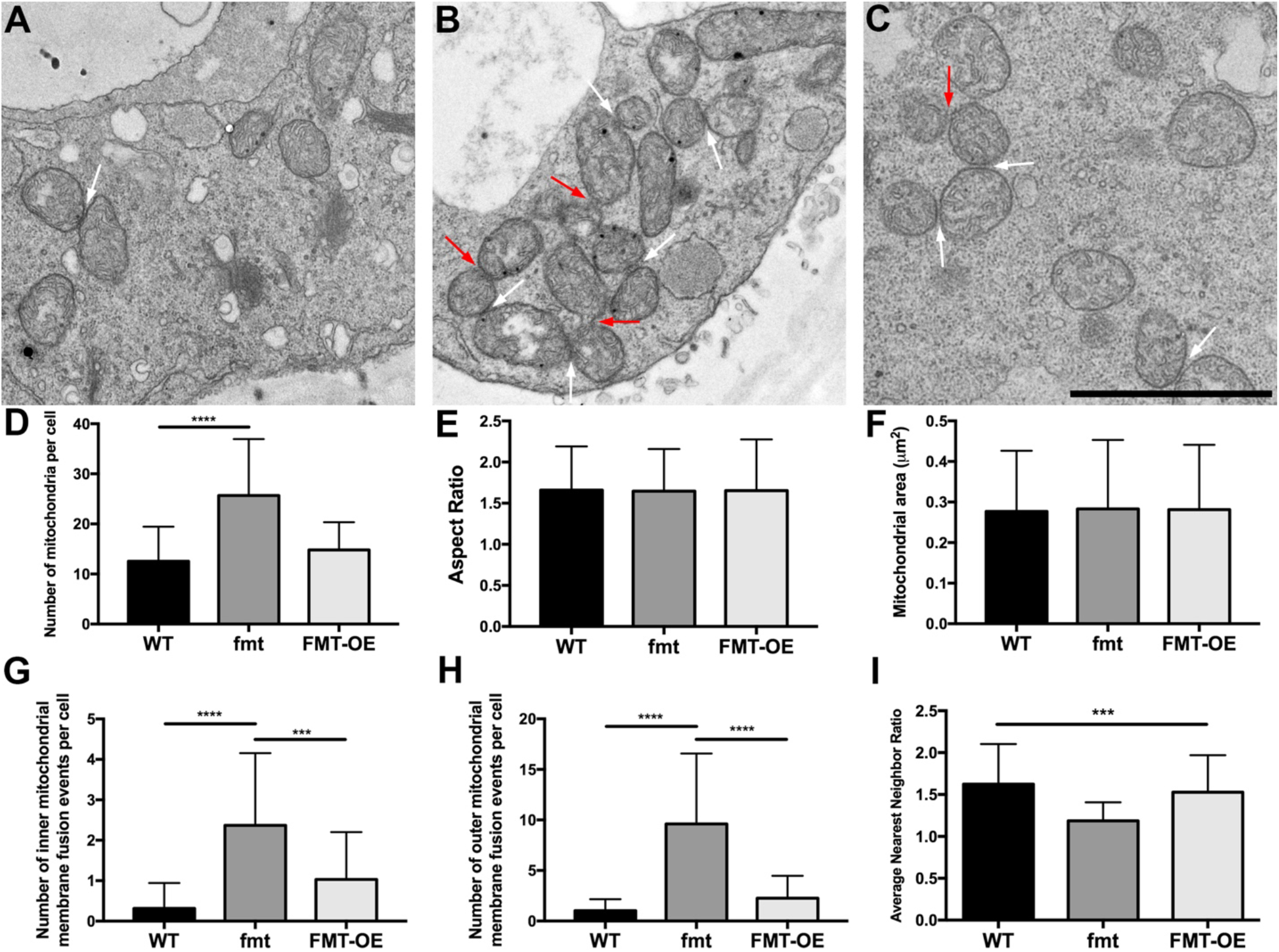
FMT regulates mitochondrial morphology and dynamics. A-C) TEM of mitochondria from columella cells. Each square represents a 4 μm x 4 μm area selected from a representative cell from each genotype: A) WT, B) *fmt* and C) *FMT-OE*. D) Number of mitochondria per cell from three plant lines. E) Aspect ratio (AR) from three plant lines. F) Mitochondrial area (μm^2^) from three plant lines. G) Inner mitochondrial membrane (IMM) fusion events from three plant lines. H) Outer mitochondrial membrane (OMM) fusion events from three plant lines. I) Nearest neighbor value (Rn) between three plant lines. Statistical analysis indicates significant differences (****, P ≤ 0.0001, ***, P ≤ 0.001, *, P ≤ 0.05) using one-way ANOVA. Bars and error bars represent mean and standard deviation, respectively. Scale bar = 2 μm

### *FMT* Affects Seedling Germination and Germination Timing

We next examined whether these changes in mitochondrial dynamics led to developmental, phenotypic, or behavioral changes in the whole plant. Seeds undergo numerous metabolic changes during development and maturation, including distinct changes in tricarboxylic acid (TCA) cycle and β-oxidation metabolites (Fait et al., 2006). Mitochondria undergo dynamic changes during seed germination, including a reactivation of metabolism and increased mitochondrial motility (Paszkiewicz et al., 2017). Because our data showed that *FMT* is highly expressed during the mid-stages of embryogenesis (Figure 1C-E, G,H), we analyzed the effects of both *FMT*-*OE* and *fmt* KO on germination. We measured germination timing and germination rate on Murashige-Skoog (MS) agar plates. We found that *FMT-OE* seedlings took between 72-96 hours to germinate, while WT and *fmt* seedlings germinated within 24-48 hours (Figure 4A,B). In addition, the germination rate of the *FMT-OE* line was significantly decreased (∼85%) compared to WT and *fmt* seeds (∼98%), which indicates that *FMT* also negatively regulates germination rate (Figure 4C). These data reveal a previously unknown role of FMT in regulating germination in *Arabidopsis*.

**Figure 4:**
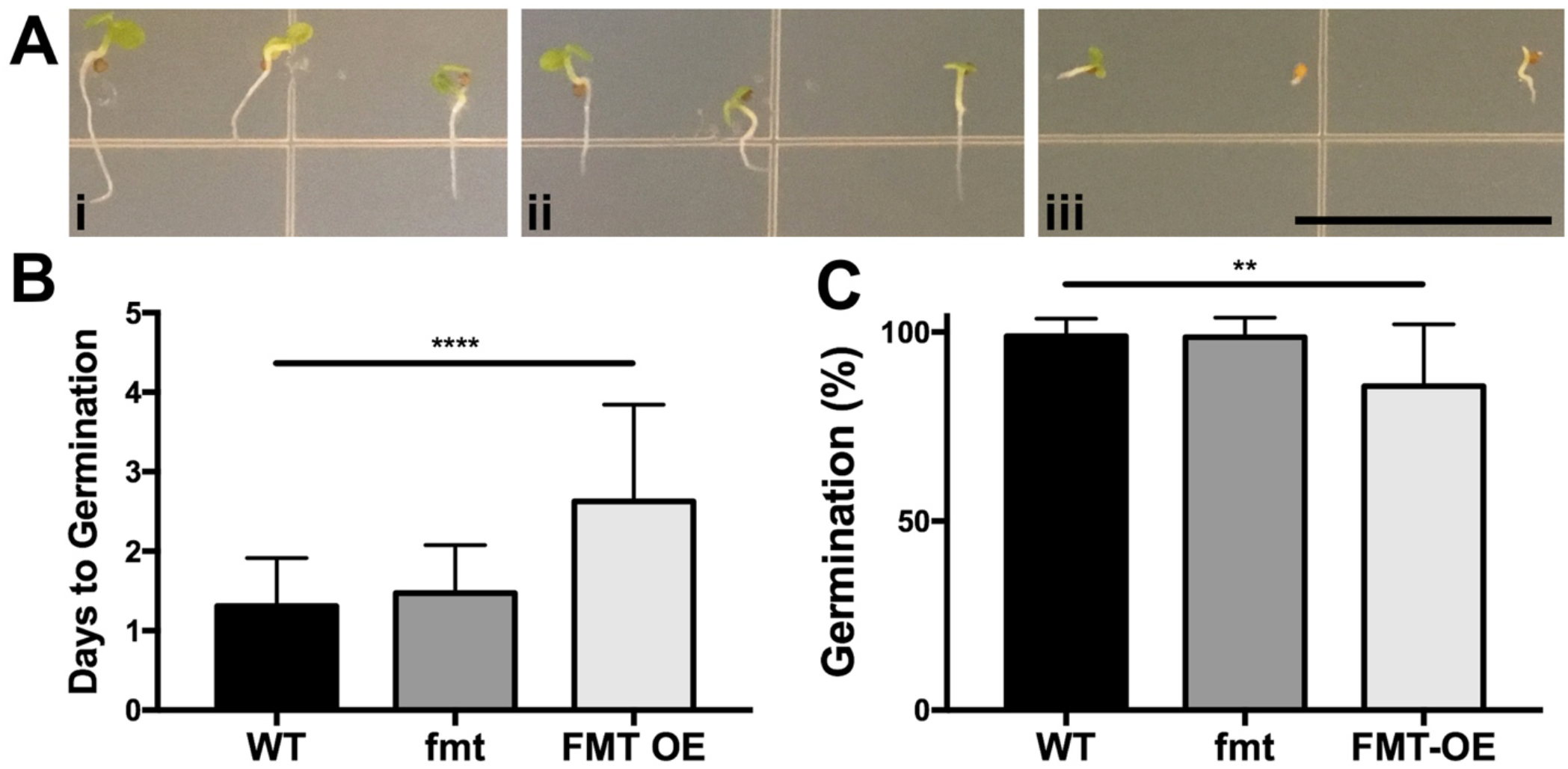
FMT regulates germination timing and total germination. A) Plants 4 days after germination (DAG), i) WT, ii) *fmt* and iii) *FMT-OE*. B) Quantification of days to germination. C) Quantification of germination rate (in %). Scale bar = 1 cm. Statistical analysis indicates significant differences (****, P ≤ 0.0001, **, P ≤ 0.01) using one-way ANOVA. Bars and error bars represent mean and standard deviation, respectively.

### *FMT* Affects Root Length and Flowering Timing in *Arabidopsis*

*fmt* mutants in *Arabidopsis* were previously described as having significantly shorter roots than the WT, likely resulting from their smaller meristematic zones (El Zawily et al., 2014). This suggests that *FMT* functions as a regulator of root length, particularly within the meristematic zone. So, we examined if *FMT* overexpression increased root length when grown on MS agar plates. At 7 days after germination (DAG), we found that both *fmt* and *FMT*-*OE* plants had significantly shorter roots than the WT (Figure 5A,B). Although the roots of *fmt* mutants were still significantly shorter than the WT at 14 DAG, the roots of *FMT-OE* mutants overcame an initial deficit in root length and were comparable to those of the WT (Figure 5C,D). These results confirm previous findings (El Zawily et al., 2014) and further demonstrate a clear role for *FMT* in regulating and maintaining root growth and length.

**Figure 5:**
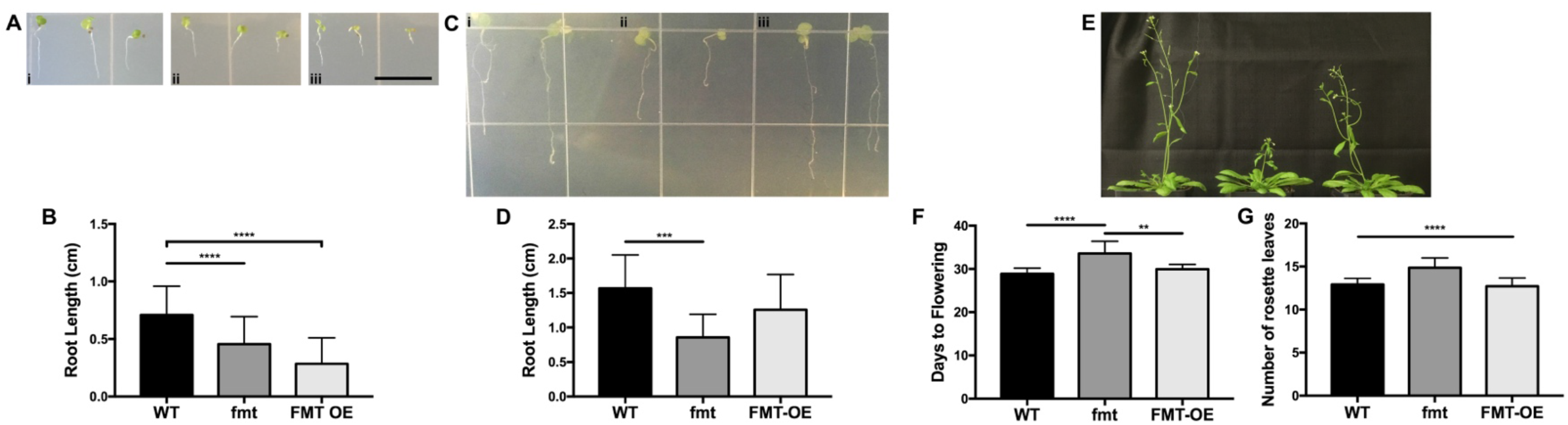
FMT regulates root length and flowering timing under long-day conditions. A) Root length at 7 DAG, i) WT, ii) *fmt* and iii *FMT-OE*. B) Quantification of root length at 7 DAG. C) Root length at 14 DAG in i) WT, ii) *fmt* and iii) *FMT-OE*. D) Quantification of root length at 14 DAG. E) Flowering time phenotype at 35 DAG, i) WT, ii) *fmt*, iii) *FMT-OE*. F) Quantification of days to flowering. Scale bar = 1 cm. Statistical analysis indicates significant differences (****, P ≤ 0.0001, ***, P ≤ 0.001, **, P ≤ 0.01) using one-way ANOVA. Bars and error bars represent mean and standard deviation, respectively.

Since we demonstrated that *FMT* is expressed in the inflorescence shoot apical meristem and the flower structure (Figure 1), we examined the effect of *FMT* on flowering timing in *Arabidopsis* under long-day (16 h light/8 h dark) conditions. We defined flowering time of WT, *fmt* and *FMT-OE* as the time point of the emergence of the first visible floral stem (bolting) and the number of rosettes leaves at the time of flowering. We found that *fmt* plants exhibited a significant delay in flowering timing, taking on average between 33-34 days to flower, compared to 28-29 days in the WT and 29-30 days in *FMT-OE* plants (Figure 5E,F). In addition, we found that *fmt* mutants had a significant increase in the number of leaves at the time of flowering compared to WT and *FMT-OE* plants (Figure 5G). Generally, the rosette leaf number reflects the timing of flowering. We found an increase in leaf number in *fmt* mutants that positively correlated with the late flowering time phenotype of the mutant. Our data indicate that *FMT* is necessary for normal flowering in the *Arabidopsis* WT, where it acts as a positive regulator of flowering timing.

### *FMT* Affects the Salt Stress Response in *Arabidopsis*

Plants exhibit several behavioral responses to salt stress, including inhibition of germination, delay in flowering timing, disrupted root growth, and salt avoidance tropism (Duan et al., 2015; Li and Zhang, 2008; Ryu et al., 2014; Sun et al., 2008; Vallejo et al., 2010). Since *FMT* regulates mitochondrial morphology and dynamics, we tested whether salt stress-responsive behaviors were altered in our mutants. Using the spatio-temporal root stress browser (Dinneny et al., 2008), we were able to examine expression levels in roots of five-day-old *Arabidopsis* Col-0 seedlings exposed to 140 mM NaCl for 48 hours. *FMT* showed an increased expression in the epidermal cell layer 3 hours after exposure to NaCl, with peak expression after 32 hours. This expression remained elevated up to 48 hours after exposure to NaCl compared to non-stressed control plants (Figure 6A). These data reveal a previously unknown role for *FMT* as a salt-responsive gene in *Arabidopsis* and indicate a major regulatory role for *FMT* in *Arabidopsis* in this physiological response. Following salt stress, plants typically decrease primary root growth to minimize root exposure to excess Na^+^ (Munns, 2002). To determine the effects of salt stress on our mutants, plants from all three lines were germinated and grown on MS agar plates for seven days, and then transferred to MS agar plates supplemented with 125 mM NaCl for seven days. We recorded root length on the seventh day. We found that both WT and *FMT-OE* plants exhibited a significant decrease in primary root length upon exposure to salt stress (Figure 6F,6H), while primary root length in *fmt* mutants remained unchanged (Figure 6G). Taken together, these data provide evidence that *FMT* regulates salt stress-induced phenotypes, including germination, flowering timing, and primary root growth.

**Figure 6:**
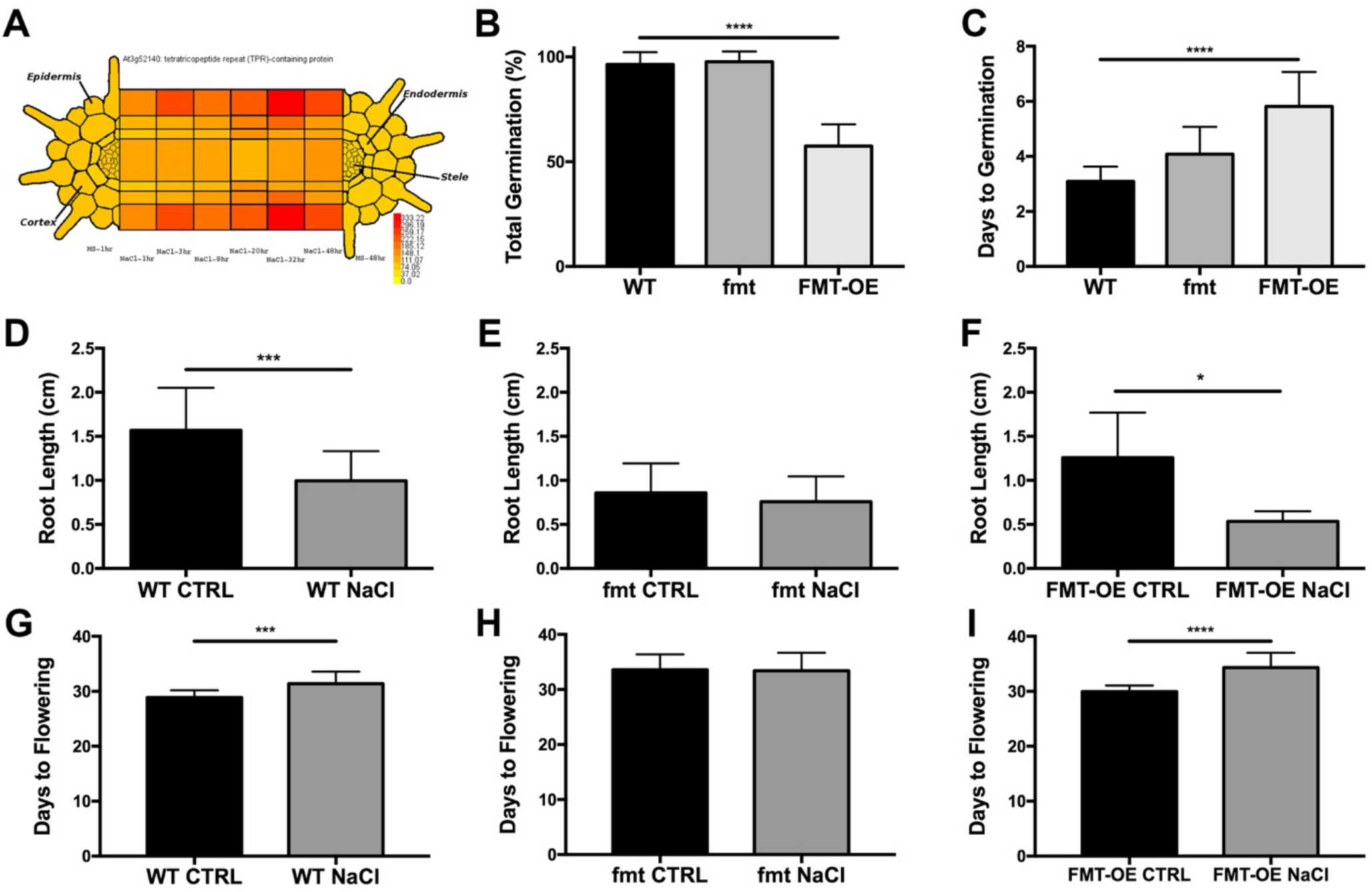
FMT affects the response to salt stress in *Arabidopsis*. A) Spatio-temporal root stress browser of *FMT* expression in response to 140 mM NaCl over 48 hours. Control (MS) expression at 1 and 48 hours provided on the left and right root cross-sections, respectively. Gene expression level is directly compared to the highest signal record for the given gene (*FMT*) and assigned an accompanying color, ranging from yellow (no expression) to red (highest/absolute) expression. B) Quantification of total germination (in %). C) Days to germination. D) Primary root length under control and salt-stress conditions in WT. E) Primary root length under control and salt-stress conditions in *fmt*. F) Primary root length under control and salt-stress conditions in *FMT-OE*. G) Days to flowering under control and salt-stress conditions in WT. H) Days to flowering under control and salt-stress conditions in *fmt*. I) Days to flowering under control and salt-stress conditions in *FMT-OE*. Black bars indicate control conditions. Gray bars indicate exposure to 125 mM NaCl. Statistical analysis indicates significant differences (****, P ≤ 0.0001, ***, P ≤ 0.001, *, P ≤ 0.05) using one-way ANOVA. Bars and error bars represent mean and standard deviation, respectively.

Excess salt in the soil is first sensed by the root cap, which transmits signals to the rest of the plant to induce salt-responsive behaviors (Deinlein et al., 2014). We examined whether altered mitochondrial dynamics in the root cap could explain these differences in behavior, since *FMT* is expressed in the root and up-regulated during salt stress, and the mutants exhibited an altered response to salt stress. Using TEM, we analyzed mitochondria in root cap cells of our three lines after exposure to 125 mM NaCl stress. We found that the number of mitochondria per cell increased significantly in WT and *FMT-OE* plants upon salt stress, but remained unchanged in *fmt* (Figure 7D-F). The number of IMM contacts decreased significantly in *FMT-OE* plants during salinity stress, but was not affected in WT or *fmt* (Figure 7G-I). In addition, OMM fusion events were not affected by salt stress in any of the three lines (Figure 7J-L). After salt stress, mitochondria trended towards dispersion in the WT, clustering in *FMT-OE* plants, while *fmt* remained unchanged compared to WT (Figure 7M-O). Total mitochondrial area remained unchanged in *fmt* mutants following salt exposure (Figure S4,A-C), while mitochondrial aspect ratio decreased significantly in all three lines (Figure S4,D-F). These results indicate that *FMT* regulates various aspects of mitochondrial dynamics and morphology under salt stress conditions. Altering this regulation could induce changes in the salt stress response of the mutants.

**Figure 7:**
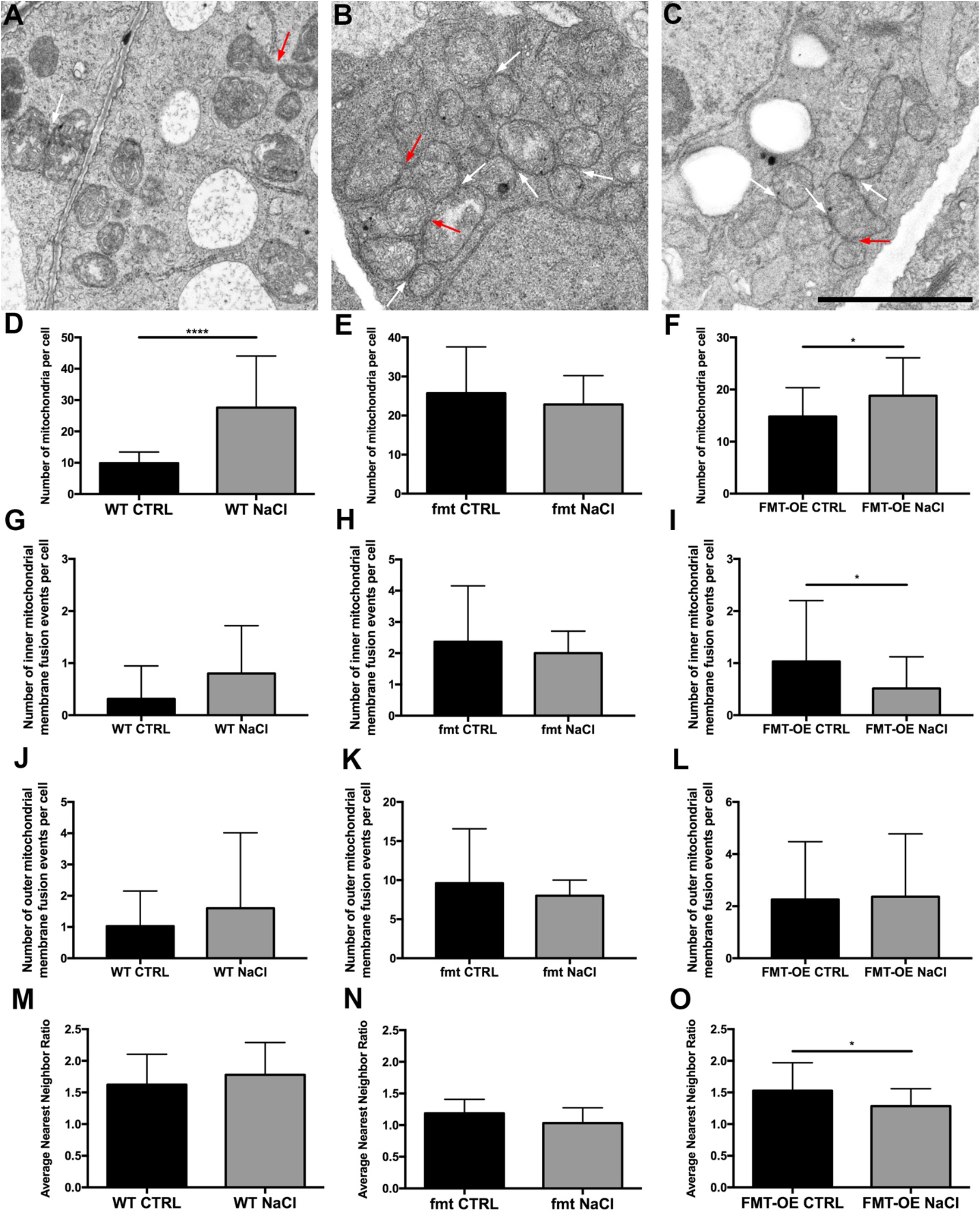
FMT regulates mitochondrial morphology and dynamics under salt stress. A-C) TEM of mitochondria from columella cells. Each square represents a 4 μm x 4 μm area selected from a representative cell from each genotype: A) WT, B) *fmt* and C) *FMT-OE*. D-F) Number of mitochondria per cell in plants under salt stress compared to control conditions in WT, *fmt*, and *FMT-OE*, respectively. G-I) Number of inner mitochondrial membrane fusion events in plants under salt stress compared to control conditions in WT, *fmt*, and *FMT-OE*, respectively. J-L) Number of outer mitochondrial membrane fusion events in plants under salt stress compared to control conditions in WT, *fmt*, and *FMT-OE*, respectively. M-O) Average nearest neighbor ratio in in plants under salt stress compared to control conditions in WT, *fmt*, and *FMT-OE*, respectively. Statistical analysis indicates significant differences (****, P ≤ 0.0001, *, P ≤ 0.05) using one-way ANOVA. Bars and error bars represent mean and standard deviation, respectively.

We further tested germination rate and timing of WT, *fmt* and *FMT-OE* plants under salt stress conditions. We sowed seeds on MS agar plates supplemented with 125 mM NaCl. We found that germination rate was significantly decreased by ∼50% in *FMT*-*OE* seeds compared to WT and *fmt* (Figure 6B). Seeds of *FMT-OE* plants took significantly longer to germinate (∼6 days) than seeds from WT and *fmt* plants (Figure 6C). Of note, salt treatment affects germination rate and flowering timing of *FMT-OE* plants to a greater degree than observed under normal (non-stress) conditions (see Figure 4). The *FMT-OE* plants germinated in ∼4 days under non-stress conditions (Figure 4A), while showing delayed germination by approximately two days under salt treatment (Figure 6C). We observed a similar response for germination rate, which was affected by 50% under stress in *FMT-OE* plants (Figure 6B) compared to ∼89% under normal conditions (Figure 4C). The *fmt* mutant displayed an insensitivity to salt treatment, showing a WT-like germination rate. However, the days to germination was also significantly increased in *fmt* compared to WT plants. Total germination rate remained unchanged compared to control conditions (Figure S3). Our above results demonstrated that *FMT* acts as a negative regulator of germination under normal conditions in *Arabidopsis*. In the presence of salt treatment, we found an enhanced stress response in *FMT* overexpression plants, which further delayed seed germination compared to control conditions. These results demonstrate that salt stress influences the germination process through the *FMT* gene.

We next exposed one-week-old seedlings to 125 mM salt stress on soil and recorded the days to flowering for each of the three lines. We found that both WT (Figure 6G) and *FMT-OE* plants (Figure 6I) took significantly longer to flower under salt stress, defined as the days to emergence of the first visible floral stem, than *fmt* mutants. Notably, salt stress did not delay flowering of *fmt* mutants compared to control conditions (Figure 6H).

Salt stress can affect flowering timing depending on the salt concentration (Ryu et al., 2014; Ryu et al., 2011). So, we sought to define when the floral transition took place in WT plants grown under salt stress. We performed toluidine blue staining on longitudinal sections of apices of Col-0 plants grown under long days and harvested at 0, 1, 2 and 3 days after transfer (DAT) to salt media (Figure 8A-H). As expected, WT plants grown under control conditions initiated the floral transition faster (1 DAT) (Figure 8B) than the salt grown plants (3 DAT) (Figure 8H). To delineate the role of *FMT* during salt exposure, we analyzed the expression of *FMT* in the SAM of WT plants grown in control and salt conditions (125 mM NaCl) by RNA *in situ* hybridization using *FMT* as a probe. We analyzed longitudinal sections of apical meristems of plants grown under either normal or salt stress conditions, harvested at end of day (ED) at 0, 1 and 2 DAT to salt media (Figure 8I-N). We detected transcripts of *FMT* at a low level in the vegetative SAM of WT plants grown in both conditions at 7 DAG (0 DAT) (Figure 8I,L). Interestingly, *FMT* expression was highly induced during floral transition in the SAM and young leaves at 1 DAT in WT plants grown in control conditions (Figure 8J). *FMT* transcript abundance initially declined after shifting plants to salt media (Figure 8M) and then recovered, as we detected the *FMT* transcript level in the SAM of salt treated plants at 2 DAT (Figure 8N). These data provide evidence that salt exposure decreases *FMT* expression in the SAM. Our observed delay in flowering time of WT and *FMT-OE* plants grown in salt media may reflect decreased *FMT* expression upon exposure to salt. Thus, our data indicates that *FMT* is an essential regulator of flowering timing and a potential target of salt-affected flowering. Taken together, these data provide evidence that *FMT* regulates salt stress-induced phenotypes, including germination, flowering timing, and primary root growth.

**Figure 8:**
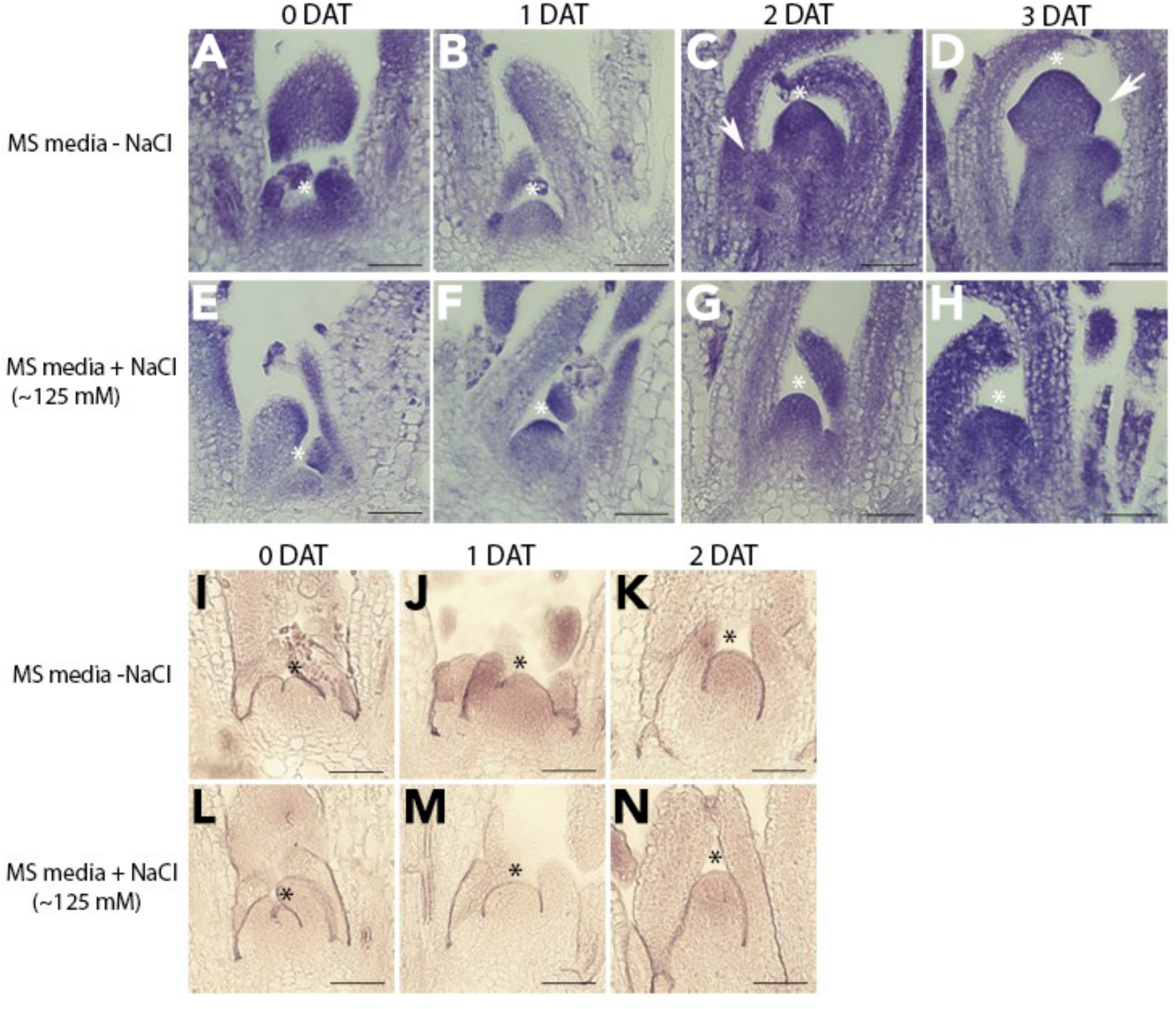
Salt treatment delays floral transition in Col-0 plants and affects *FMT* expression in the shoot apical meristem. A-H) Emergence of flower primordia analyzed by toluidine blue staining from apices of Col-0 plants grown under long days and harvested at 0, 1, 2 and 3 days after transfer (DAT) to salt media. I-N) RNA *in situ* hybridization using *FMT* as a probe on longitudinal sections of the apical meristem of shifted plants grown in normal and salt conditions, harvested at end of day (ED) at 0, 1 and 2 days after transfer (DAT). Asterisks and arrows indicate meristem summit and floral primordia, respectively. Scale bars = 100 μm.

### *FMT* Affects Hyponasty and Plant Size by Regulating Leaf Expansion Growth

Since *fmt* mutants show a reduced plant size (El Zawily et al., 2014), we tested whether *FMT* regulates expansion growth using an established phenotyping system (Phytotyping^4D^) (Apelt et al., 2015). We imaged eight WT, ten *fmt*, and eight *FMT-OE* plants using the Phytotyping^4D^ system for eight consecutive days (from 15 to 23 days after sowing, DAS) at a time resolution of five images per hour per plant (Figure 9A). During this time, all three genotypes showed an exponential increase in rosette area, but *fmt* plants were approximately 25% smaller (p<0.05, Student’s *t*-test), and *FMT-OE* were approximately 35% smaller (p<0.01, Student’s *t*-test) compared to the WT at 23 DAS. The time series from the eight 24-hour-cycles were combined to calculate the diurnal changes in the relative expansion rate (RER) (Figure 9B,C). Generally, the diurnal RER of the three lines resembled those reported previously (Apelt et al., 2015). RER decreased after dawn, peaked around 4 h, and then stayed fairly constant for the remainder of the day with a peak directly after dusk. We found only marginal diurnal differences among the three lines. For example, there were slight variations in RER timing, as *fmt* mutants had a significantly enhanced RER peak at 4 hours after dawn compared to WT and *FMT-OE*.

**Figure 9:**
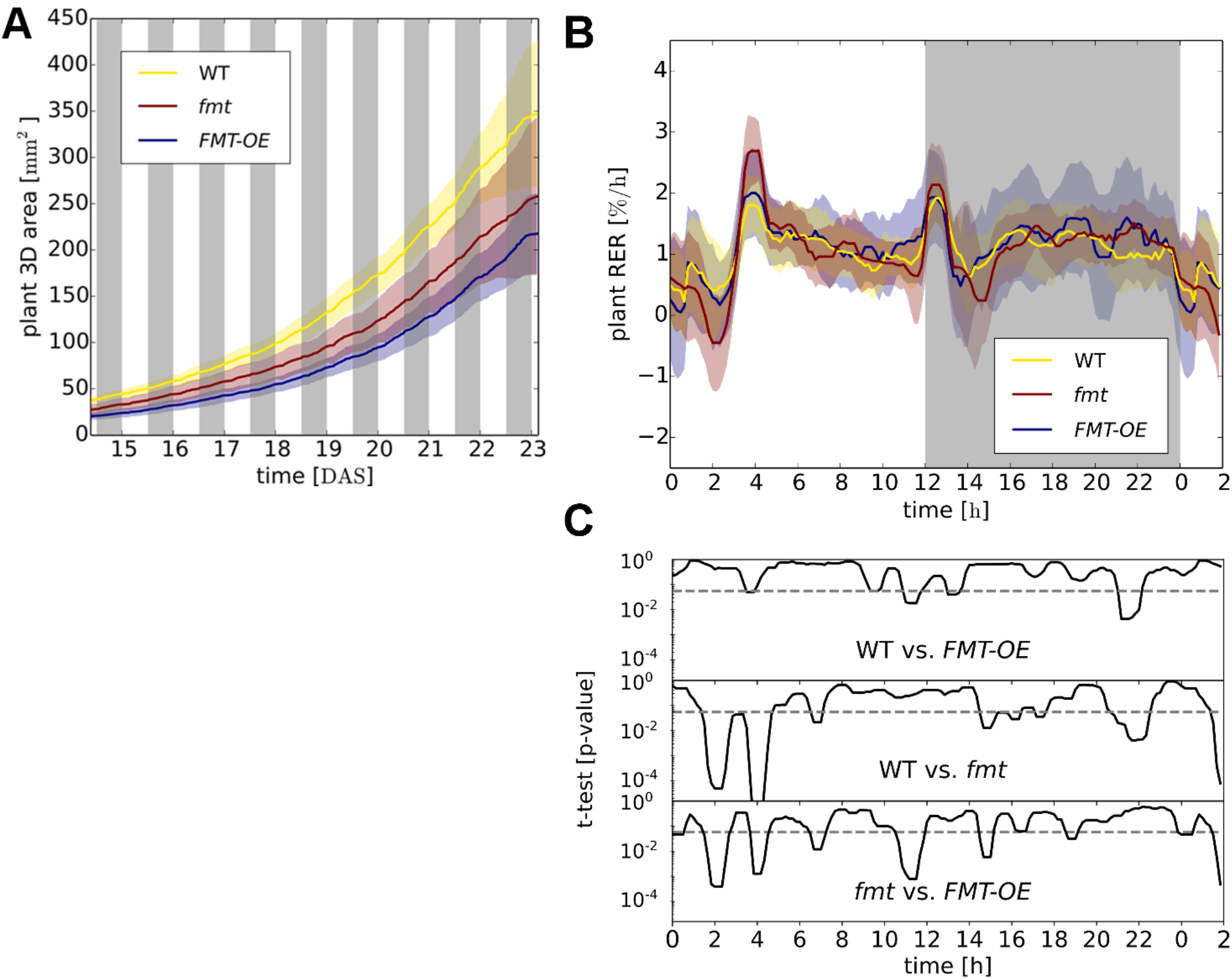
*fmt* and *FMT-OE* mutants exhibit changes in spatio-temporal growth. A) Rosette area increased over time (15-23 days after sowing, DAS) determined by 3D imaging. Yellow: WT, red: *fmt*, blue: *FMT-OE*. B) Diurnal relative expansion rate (RER) averaged over eight sequential 24-hour periods. C) p-values from pairwise Student’s *t*-tests applied over a 50 minute sliding window, where p-values <0.05 indicated significance. Lines and color-shaded areas represent mean and standard deviation, respectively. Plants grown in a 12 h photoperiod.

Behavior has been defined as: “what a plant or animal does, in the course of an individual’s lifetime, in response to some event or change in its environment” (Silvertown and Gordon, 1989). This definition is not intended to assume that all behaviors exhibited by animals are also present in plants and *vice versa*; rather, it comprises a more comprehensive definition to encompass the range of overlap between the behaviors of plants and animals in response to their environments. We can classify these behaviors based on various activities, such as growth (of an individual plant or animal), irreversible change (seed germination in plants or imprinting in animals), flexible change (movements based on circadian clocks in both plants (nyctinasty/hyponasty) and animals (sleep/wake cycle)), or response to stress (avoidance in both plants and animals).

Because *fmt* mutants exhibit altered mitochondrial dynamics and distribution that elicits phenotypic and behavioral changes in animals, we analyzed behavioral changes in *OE* and mutant plants. We evaluated behavior of our three plant lines by examining movements, i.e. mobility based on the circadian clock, and stress-responsive behaviors, which can be accurately detected and measured. Hyponastic movement of *Arabidopsis* leaves follows a pattern of rhythmic, diurnal oscillation driven by the circadian clock (McClung, 2006), which is measured as the change in leaf angle over time (Muller and Jimenez-Gomez, 2016). We employed the Phytotyping^4D^ system to determine the hyponastic behavior of leaves by tracking their average hyponastic angles from 15 to 23 DAS. We found that WT plants exhibited a high leaf angle during the night, which decreased after dawn, and then increased at dusk and throughout the night. These results are consistent with previous reports from Apelt et al. (2015) and Apelt et al. (2017). In our tested transgenic and mutant lines, we saw a consistently higher hyponastic angle in *FMT-OE* plants, while *fmt* mutants displayed a consistently lower hyponastic angle (Figure 10A). This finding was even more pronounced when incorporating the diurnal average of the hyponastic angle over the whole time series (Figure 10B,C). *fmt* mutants showed a temporal delay of leaf angle increase after dusk and lower angles during the night. We also found significant differences in the speed of leaf movement between the fmt mutant and WT and *FMT-OE* plants after dusk and during the first half of the day (Figure 10D,E).These data demonstrate an important role for *FMT* in regulating hyponastic behavior, including changes in speed of hyponastic movements.

**Figure 10:**
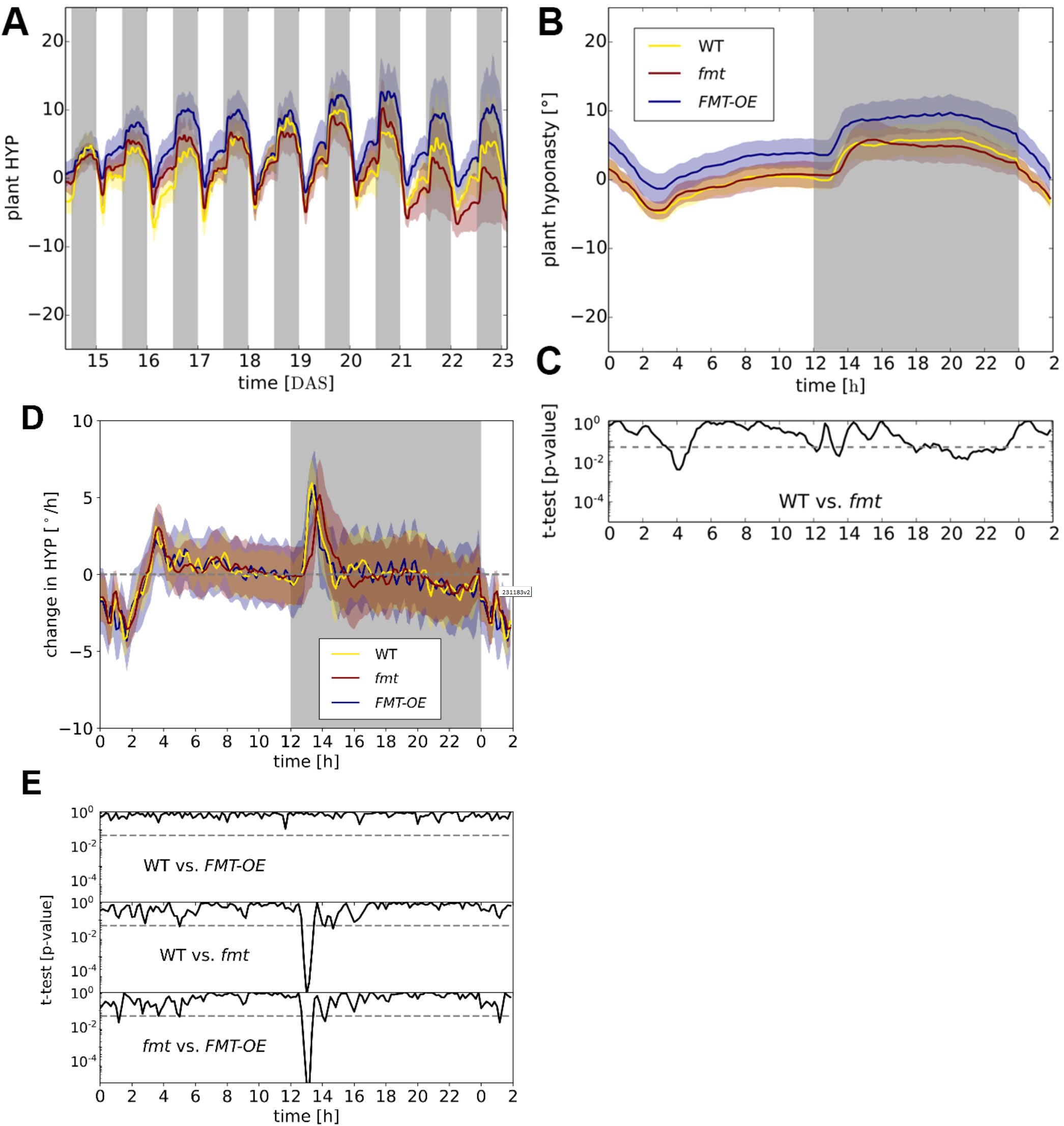
FMT regulates hyponastic growth in *Arabidopsis*. A) Average hyponastic angle (HYP) over time (15-23 DAS). B) Diurnal HYP averaged over eight sequential 24-hour periods. C,E) p-values from pairwise Student’s *t*-tests applied over a 50 minute sliding window, where p-values <0.05 indicated significance. D) Change in speed of hyponastic angle (HYP) averaged over eight sequential 24-hour periods. Lines and color-shaded areas represent mean and standard deviation, respectively. Plants grown in a 12 h photoperiod.

### Knockout of CLUH in Mice Affects Mobility and Speed

The homologue of FMT in mice, CLUH (clustered mitochondria homologue), has previously been shown to bind nuclear-encoded mitochondrial mRNAs (Gao et al., 2014). However, the effects of CLUH on behavior in mice has yet to be reported. To examine whether CLUH may have behavioral effect in mice corresponding to the effect of *fmt* in *Arabidopsis*, we studied whole body heterozygous (HET) *Cluh*^+/-^ mice (Schatton et al., 2017). There were no observable differences in bodyweight between WT and *Cluh*^+/-^ mice of either sex at 8 weeks of age (Figure S5). We performed several behavioral tests, including marble burying, open field, and zero maze on 18 WT and 17 HET mice. We found no significant differences in the marble burying test or zero maze test (data not shown). However, we observed significant differences between WT and heterozygous mice in the open field test. We found that heterozygous mice travelled for shorter distances (Figure 11A), and had a slower average and maximum speed (Figures 11B and 11C, respectively). These results unmask a conserved and overlapping influence of *fmt* and CLUH on speed of movement of plants and animals, respectively.

**Figure 11:**
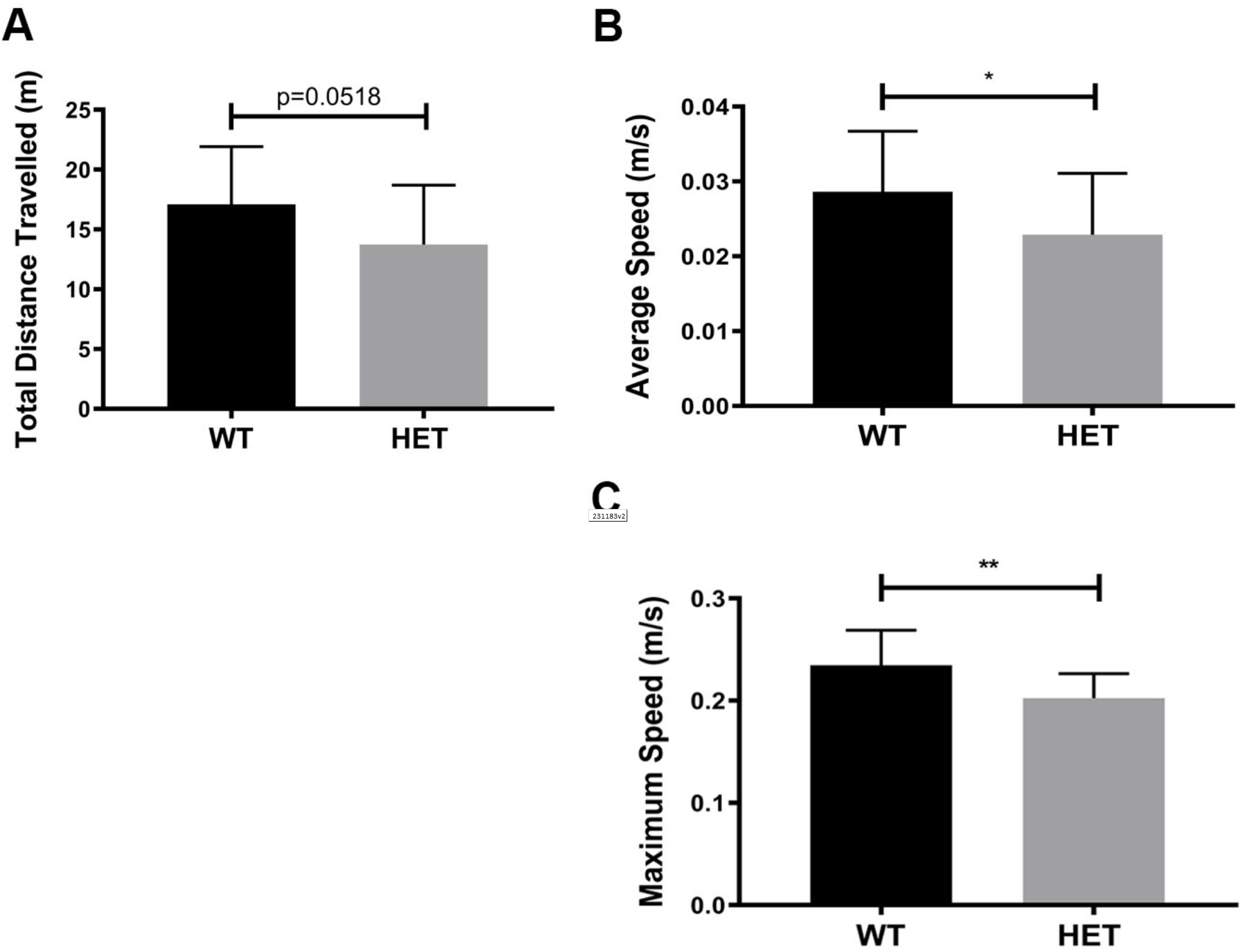
CLUH affects mouse mobility and speed. A) Comparison of WT (n=18) and heterozygous (HET) *Cluh*^+/-^ mice (n=17) in open field tests: A) Total distance travelled, B) Average speed, C) Maximum speed.

## DISCUSSION

Mitochondria are highly conserved organelles essential for the health of both plants and animals. They regulate numerous critical processes, such as energy production, cellular metabolism, and apoptosis, among others (Chan, 2006; Cheng et al., 2010; Mattson et al., 2008). Mitochondrial dysfunction is implicated in underlying multiple physiological and pathological behaviors in higher organisms and hallmark behaviors of multiple neurodegenerative diseases, including Alzheimer’s disease, Parkinson’s disease, Schizophrenia, Williams syndrome and autism spectrum disorders (Bertholet et al., 2016; Burte et al., 2015; Golpich et al., 2017; Goncalves et al., 2015; Islam, 2017; Krols et al., 2016; Oka, 2016; Rajasekaran et al., 2015). Plants and animals share numerous homologous genes encoding mitochondrial proteins that are in themselves highly functionally conserved, including *CLU*, a gene originally thought to mediate mitochondrial localization (Fields et al., 1998; Zhu et al., 1997). However, we now know that CLU plays multiple important roles within the cell, including regulating mitochondrial autophagy through the interaction between PINK1 and Parkin (Sen et al., 2015) and mitochondrial metabolism (Schatton et al., 2017) and directing mitochondrial trafficking (El Zawily et al., 2014). *FMT*, the functional homolog of *CLU* in *Arabidopsis*, regulates not only various aspects of mitochondrial morphology and dynamics within the cell, as well as plant development (El Zawily et al., 2014), but also hyponastic and salt stress-responsive behaviors.

We confirmed previous findings that *fmt* mutants exhibit altered mitochondrial morphology and dynamics, hence *FMT-*deficient plants show a significantly increased number of mitochondria and display more clustered mitochondria (El Zawily et al., 2014). Mitochondria of *fmt* mutants also show more connections with each other, either through inner or outer mitochondrial membrane contacts. However, we found also that under salt stress conditions, the number of mitochondria per cell increased significantly in WT and *FMT-OE* plants but remained unchanged in *fmt* mutants. These results reveal a vital role of *FMT* in regulating mitochondrial morphology and dynamics under both control and salt stress conditions.

Behavior is a fundamental principle that occurs when an organism responds to its environment. It follows a distinct hierarchy that begins with changes in cellular and molecular mechanisms that then drive changes in various physiological processes through which phenotypic and developmental changes emerge. Understanding these cellular and molecular driving forces will enable a greater understanding of the fundamental behaviors themselves. If the underlying behavioral processes are conserved and universal, we propose that fundamental cellular and molecular mechanisms drive behavior. These mechanisms then should themselves be conserved among organisms and species, such as plants and animals. Studying the cellular and molecular mechanisms that drive behavior, including those that are highly functionally conserved, will permit a detailed examination of the fundamental principles that underlie physiological and pathological states in multiple species. We found that alterations in the mitochondrial dynamics of our mutants altered both hyponastic and salt stress-response behaviors. Our results suggest that further work using our proposed perspective will elucidate the mechanisms driving these changes to reveal fundamental principles inducing phenotypic and behavioral changes in higher organisms, including animals.

## AUTHOR CONTRIBUTIONS

Conceived and designed study, A.R. and T.L.H.; performed experiments, A.R., F.A., and J.J.O.; analyzed results, A.R., F.A., and J.J.O.; provided essential materials, F.K., B.M.R., E.I.R. and T.L.H.; wrote the manuscript, A.R. and T.L.H; edited manuscript, all authors.

## ACKNOWLEDGMENTS

This work was supported by AG052005, AG052986, AG051459, DK111178 from NIH (T.L.H). Work in the Mueller-Roeber group is supported by the Deutsche Forschungsgemeinschaft (DFG) grant within the Collaborative Research Centre 973 ‘Priming and Memory of Organismic Responses to Stress’ (www.sfb973.de). We thank Life Science Editors for editing assistance.

## SUPPLEMENTAL FIGURES

**Figure S1 (related to Figure 2). Agarose gel of PCR results of WT and *fmt* knockout lines**. A) DNA size marker. B) WT with primers spanning the insertion site. C) WT with primers including the T-DNA border site. D) *fmt* with primers spanning the insertion site. E) *fmt* with primers including the T-DNA border site.

**Figure S2 (related to Figure 2). *FMT-OE* exhibits a significant increase in mRNA expression of *FMT***. Quantification of *FMT* expression in *fmt* and *FMT-OE* lines. *FMT-OE* plants exhibit an almost two-fold higher mRNA expression of *FMT*, while there is no detectable expression in *fmt* plants. Statistical analysis indicates significant differences (**, P ≤ 0.01) using one-way ANOVA. Bars and error bars represent mean and standard deviation, respectively.

**Figure S3 (related to Figure 6). Salt stress delays time to germination but not total germination in WT and *fmt***. A) Quantification of days to germination in WT. B) Quantification of days to germination in *fmt*. C) Quantification of germination rate (in %) in WT. D) Quantification of germination rate (in %) in *fmt*. Statistical analysis indicates significant differences (****, P ≤ 0.0001) using one-way ANOVA. Bars and error bars represent mean and standard deviation, respectively.

**Figure S4 (related to Figure 7). Comparison of aspect ratio and mitochondrial area under control and salt stress conditions**. A-C) Aspect ratio of mitochondria in plants under salt stress compared to control conditions in WT, *fmt*, and *FMT-OE*, respectively. D-F) Mitochondrial area in plants under salt stress compared to control conditions in WT, *fmt*, and *FMT-OE*, respectively.

**Figure S5 (related to Figure 11). Bodyweights of WT and *Cluh***^**+/-**^ **mice at 8 weeks of age**. A) Comparison of bodyweights between male WT and *Cluh*^+/-^ mice. B) Comparison of bodyweights between female WT and *Cluh*^+/-^ mice.

## STAR * METHODS

### Plant materials and growth conditions

All experiments were done using Arabidopsis (*Arabidopsis thaliana*) in the Columbia-0 (Col-0) ecotype. Seeds were surface sterilized by washing in 70% (v/v) ethanol for 5 s, and this wash was replaced by 0.1% triton X-100 in 50% household bleach (v/v) for 5 sec before five rinses in sterile distilled water. Seeds were then plated on 100 × 100 × 15 mm square petri dishes (Ted Pella) filled with MS-agar, and plates were stratified at 4°C for three days in the dark to synchronize germination. Plates were then moved to long-day photoperiod conditions at 22°C +/-1°C under cool-white light at 100 μmol m^-2^ s^-1^. Plates were placed at an angle to allow for root growth along the surface of the agar. For salt-stressed growth conditions on plates, seeds were plated on MS-agar plates supplemented with 125 mM NaCl. For experiments done in soil, Arabidopsis seeds were stratified before direct sowing on Fafard #2 soil mixture (Sun Gro Horticulture)

### T-DNA insertions lines

*FMT* was screened for available T-DNA insertion lines on TAIR (The Arabidopsis Information Resource, http://www.arabidopsis.org/). PCR was used to test whether the T-DNA was inserted at the predicted insertion site. All T-DNA insertions were confirmed via PCR using left and right primers flanking the genomic sequence, and a border primer located within the T-DNA sequence (Table 1).

**Table 1:**
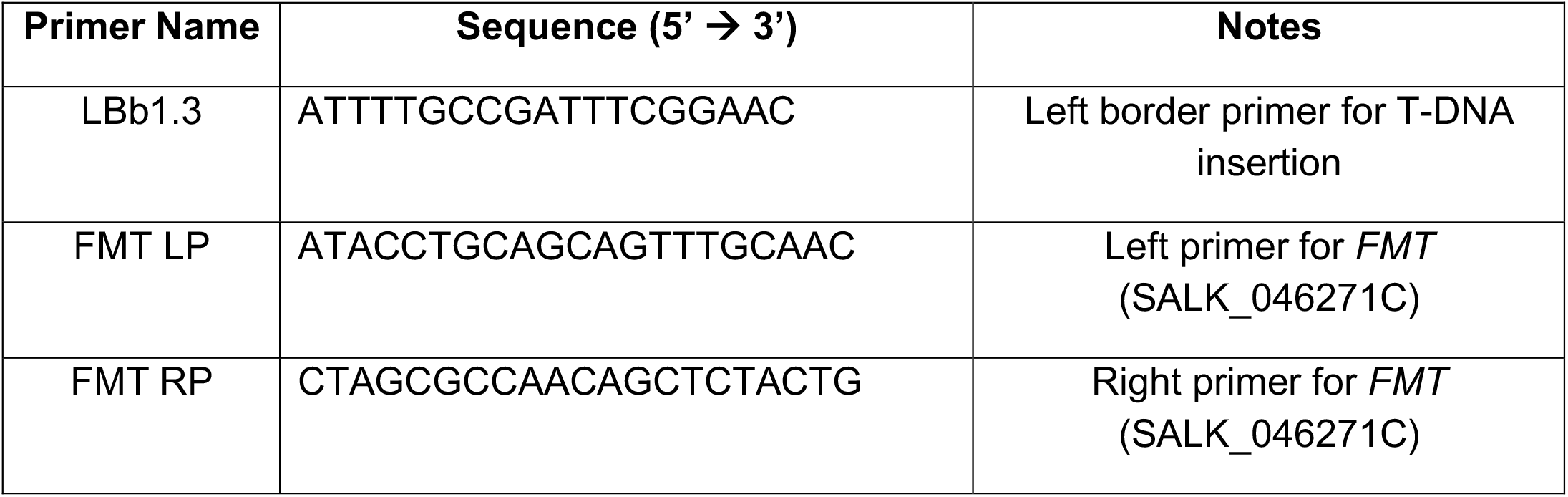
T-DNA primers used in this study.

### *FMT* cloning

pFASTG02-FMT was constructed by subcloning a *FMT* cDNA fragment of the expected size into the pDONR 221 vector (Gateway, Invitrogen). In order to PCR amplify the cDNA fragment, the following primers were used: forward (attB1FPFMT), 5’-GGGGACAAGTTTGTACAAAAAAGCAGGCTTCATGGCTGGGAAGTCGAAC-3’, and reverse (attB1RPFMT), 5’-GGGACCACTTTGTACAAGAAAGCTGGGTCTTTTTTGGCTTTTTGCTTCTT-

3’. The fragment was subsequently cloned into pFASTG02 (Shimada et al., 2010) according to the manufacturer’s protocol (Invitrogen). This vector construct, pFASTG02-FMT, allowed for overexpression of the *FMT* gene under the control of the cauliflower mosaic virus (CaMV) 35S promoter. The construct was sequenced to identify an error-free clone and subsequently transformed into wild type Col-0 plants by means of *Agrobacterium*-mediated transformation using the *Agrobacterium tumefaciens* strain GV3101 using the floral dip method with the following modifications: Silwet L-77 was added to the sucrose solution to a volume of 0.05% (v/v). The pFAST vector carries a screenable GFP marker producing a signal detectable in the mature seed coat of transformed plants. Lines with a single transgene insertion were identified by an ∼3:1 segregation ratio of GFP to no-GFP, respectively, using a Zeiss Axioplan 2 fluorescence microscope (Carl-Zeiss, Germany). Transgenic GFP seeds from these lines were sown to identify a homozygous plant (FMT-3), which was identified by T3 seed that exhibited 100% GFP fluorescence compared to WT seed. Seeds from line FMT-3 were collected and used in subsequent experiments.

For germination experiments, Arabidopsis seeds were plated on square petri dishes containing MS-agar with or without supplementation of 125 mM NaCl. Total germination was recorded 7 days after sowing. For quantification of root length under control conditions, Arabidopsis plants were grown on control MS-agar plates and roots were measured using a ruler at days seven and fourteen. For quantification of root length under salt-stressed conditions, Arabidopsis plants were grown on control MS-agar plates for one week, and then transplanted to either control or 125 mM NaCl MS-agar plates for one week and roots were measured at day fourteen. For MS-agar plate experiments, at least 20 replicates were used for each experiment, and each experiment was repeated three times.

### Flowering time analysis

For flowering timing experiments, plants were grown on soil in LD at 22°C. Flowering timing was defined as the emergence of the first visible floral stem formation (bolting), as well as the number of rosette leaves at the first visible stem formation. At least 10 replicates were used for each experiment, and each experiment was repeated three times. Mutant genotypes were compared to the wild type under both control and salt-stressed conditions. Statistical differences were determined using Student’s two tailed *t* -test, one-way analysis of variance (ANOVA), or two-way ANOVA, where appropriate.

### RNA *in situ* hybridization and histological staining

For RNA *in situ* hybridization and toluidine blue staining plants were grown on MS agar plates under LD at 22°C. Meristem (vegetative and inflorescence) and silique samples were harvested, fixed and embedded into wax using an automated tissue processor (Leica ASP200S, Leica, Wetzlar, Deutschland) and embedding system (HistoCore Arcadia, Leica). Sections of 8 μm thickness were prepared using a rotary microtome (Leica RM2255; Leica).

Briefly, for toluidine blue staining slides containing sections were incubated twice in Histoclear solution (Biozym Scientific, Hessisch Oldendorf, Germany) and an ethanol series: 100% EtOH for 2 min, 100% EtOH for 2 min, 95% of EtOH for 1 min, 90% of EtOH for 1 min, 80% of EtOH for 1 min, 60% of EtOH + 0.75% of NaCl for 1 min, 30% EtOH + 0.75% of NaCl for 1 min, 0.75% NaCl for 1 min and PBS for 1 min. Afterwards, the slides were left to dry at 42°C and then incubated in 0.01% toluidine blue/sodium borate solution for 2 min, briefly washed with water and 80% EtOH. The sections were imaged with the Nikon eclipse E600 microscope using NIS-Elements BR 4.51.00 software. Adobe Photoshop CS5 was used to generate the figures.

RNA *in situ* hybridizations were performed by processing slides containing sections through the Histoclear and ethanol series. Next, slides were incubated in Proteinase K (Roche, Mannheim, Germany) and *FRIENDLY MITOCHONDRIA* (*FTM)* probes were mixed with hybridization buffer and applied to the slides for overnight hybridization. The *FMT* probes were amplified and cloned into pGEM-T Easy Vector (Promega, Madison, Wisconsin, USA) using the following primers: forward (FMTpGEMFW) 5’-ATGGCTGGGAAGTCGAAC-3’ and reverse (FMTpGEMRV) 5’-TTATTTTTTGGCTTTTTGCTTCTTC-3’, and synthesized with the DIG RNA Labeling Kit (Roche). After the hybridization overnight, slides were washed out and incubated for with 1% blocking reagent (Roche) in 1 × TBS/0.1% Triton X-100. For immunological detection, the Anti-DIG antibody (Roche) solution diluted 1:1,250 in blocking reagent was applied to the slides. For the colorimetric detection, the NBT/BCIP stock solution (Roche) diluted 1:50 in 10% polyvinyl alcohol (PVA) in TNM-50 was applied to the slides. The slides were incubated overnight and sections were imaged as described above.

### Transmission electron microscopy (TEM)

Arabidopsis seedlings were grown on MS-agar plates for one week, and were then transferred to either 125 mM NaCl MS-agar plates or MS-agar plates without the addition of NaCl for an additional week. Fixation and embedding of 14-day-old root samples was done according to (Wu et al., 2012) with the following modifications: Durcupan epoxy resin (Sigma) was used for infiltration, tissue was collected on single slot copper grids (EMS) coated with formvar, and no post-sectioning heavy metal staining was used. Transverse sections were cut ∼30 μm deep into the columella of the root and subsequently viewed under a Tecnai 12 Transmission Electron Microscope (FEI, USA). At least ten cells from four biological samples each of WT, *fmt*, and *FMT*-*OE* roots were examined for control experiments, and at least ten cells from three biological samples each of WT, *fmt*, and *FMT*-*OE* roots were examined for salt-stressed experiments.

### Cell quantification and analysis

Using Fiji, individual cells, nuclei, vacuoles, and mitochondria were traced and measured to give the following parameters: area, aspect ratio (AR), number of mitochondria per cell, and centroid XY coordinates of each individual mitochondrion. Mitochondrial coverage was calculated as a percent using the following formula: ((cytoplasmic area–mitochondrial area)*100), where cytoplasmic area = (cell area–nuclear area–vacuole area), and mitochondrial area is the sum of all the areas of the individual mitochondria within the cell. Mitochondrial clustering was calculated using the Nearest Neighbor Distances (NND) tool in the BioVoxxel toolbox plugin in Fiji (http://imagej.net/BioVoxxel_Toolbox). The NND tool measures the average nearest neighbor ratio (ANN), which is given as:

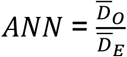

where 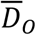 is the observed mean distance between each feature and its nearest neighbor:

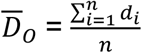

and 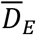 is the expected mean distance for the features given in a random pattern:

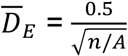

In the above equations, *d*_*i*_ equals the distance between feature *i* and its nearest neighboring feature, *n* corresponds to the total number of features, and *A* is the area of a minimum enclosing rectangle around all features.

The average nearest neighbor z-score for the statistic is calculated as:

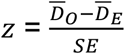

where:

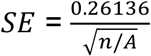

If the average distance is less than the average of a hypothetical random distribution, the mitochondria are considered clustered. If the average distance is greater than a hypothetical random distribution, the mitochondria are considered dispersed.

### Experimental set-up for measuring 3D growth behavior using Phytotyping^4D^

The Phytotyping^4D^ system provides continuous 3D recordings using a light-field camera, which allows the viewer to very accurately measure the growth rate of a plant. In particular, the camera provides both a 2D focus and a depth image simultaneously, allowing for a 3D surface reconstruction of the plant (Apelt et al., 2015; Apelt et al., 2017). Furthermore, by employing near-infrared light, Phytotyping^4D^ can continuously and non-invasively record growth over several days during light and dark periods. Arabidopsis plants were grown in 10 cm diameter pots that were filled with soil and watered containing fungicide and boric acid solution. In each pot, four equidistant spots were sown with 10–20 seeds per spot and kept for 3 days covered from light in a cold room (4°C) for stratification. Next, the plants were grown in 12 h light /12 h dark neutral days in a growth chamber (model E-36L; Percival Scientific Inc., http://www.percival-scientific.com/) with 20°C during the day and 18°C in the night and a photosynthetically active radiation of 160 μmol m^-2^ s^-1^ at the plant level. After one week, plants were thinned to one plant per spot and 15 days later the imaging was performed in the same growth chamber with Phytotyping^4D^ for 8 days. To reduce near-infrared light reflection, the soil surface was covered with black plastic-coated quartz sand before imaging. Recordings were analyzed as described in Apelt et al. (2015) obtaining non-invasive measurements of growth, RER, and hyponasty with high spatio-temporal resolution.

### Behavioral assessments of *Cluh*^+/-^ mice

The Institutional Animal Care and Use Committee of Yale University approved all experiments. Mice were kept under standard laboratory conditions with free access to standard chow food and water except during behavioral testing periods. The generation of *Cluh*^+/-^ mice has been previously described (Schatton et al., 2017). Animals included in behavioral studies were between 8 weeks of age. A series of behavioral assessments were used to establish the behavioral phenotype of *Cluh*^+/-^ mice, including marble bury, open field, and elevated zero maze. All behavioral testing was done in the light phase.

#### Marble burying test

The marble-burying test was as described (Deacon, 2006) with the following modifications: The test was performed in a static mouse cage containing 24 evenly distributed marbles arranged in a 4 x 6 pattern with alternating blue and white marbles. The marbles were placed on top of 5 cm of mouse bedding.

#### Open field

The open field apparatus is a square, polyurethane box (35.5cm x 35.5cm x 30cm). The animal was placed in the center of the apparatus. General locomotion parameters (distance traveled, locomotion speed, time mobile) and parameters relating to anxiety (freezing time; time spent, distance traveled and entries into central and periphery zones) were recorded for 10 min. The apparatus was cleaned with 70% ethanol after each animal exposure. ANY-Maze Software (Stoelting Company, Wood Dale, IL) was used to record and analyze the behavioral data.

#### Elevated zero maze

The elevated zero maze is an elevated (60cm high) ring-shaped runway (5cm wide), with 2 equally sized (25% of the runway length) sections closed off by walls (40cm high) opposite each other. The other two sections are open. The maze is equally illuminated on all four sections. Mice were placed on the center of one of the open sections, facing one of the closed sections, and allowed to explore the maze for 5 min. The apparatus was cleaned with 70% ethanol after each animal exposure. ANY-Maze Software (Stoelting Company, Wood Dale, IL) was used to record and analyze the behavioral data.

