## Supplemental Figures for "The mammalian CLU homolog *FMT* controls development and behavior in *Arabidopsis*" (from another version of this preprint: version 1, 2020-08-03)

### Supplemental Figure S1

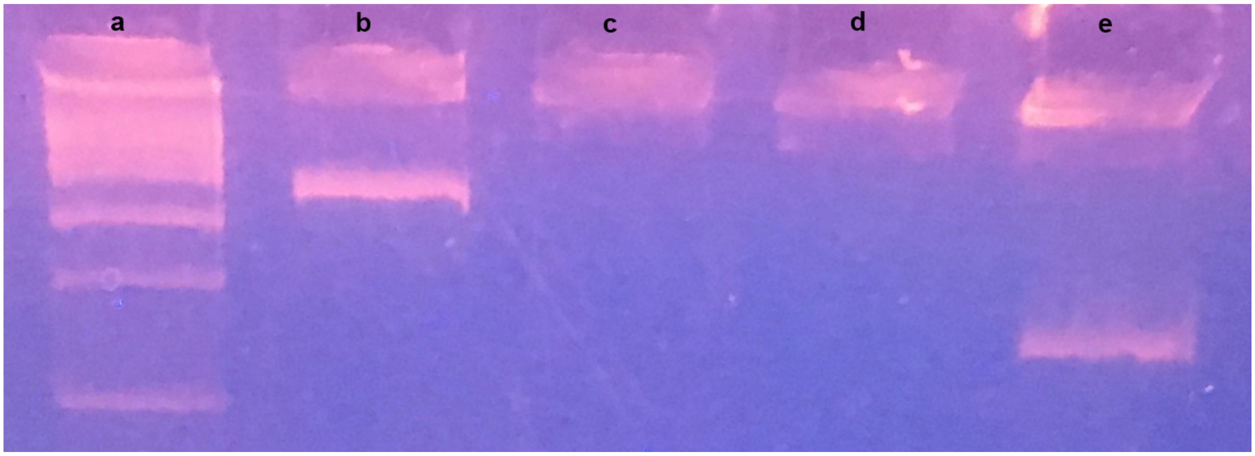


### Supplemental Figure S2

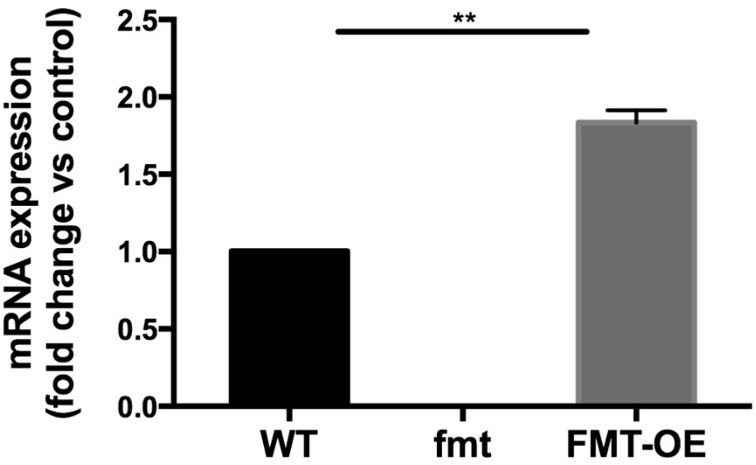

### Supplemental Figure S3

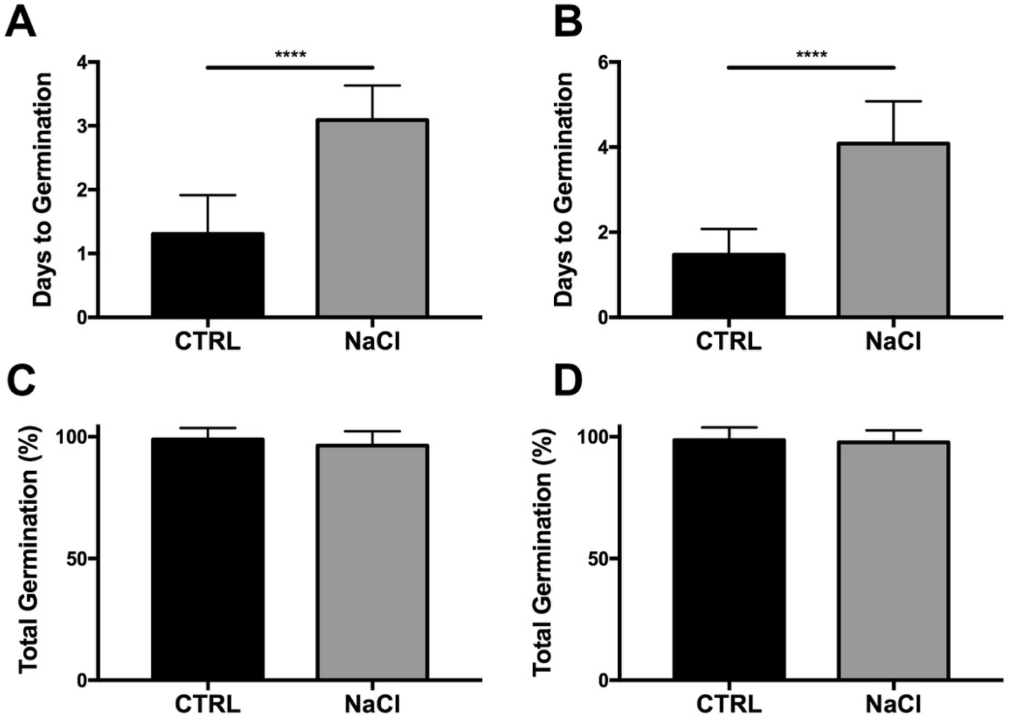


### Supplemental Figure S4

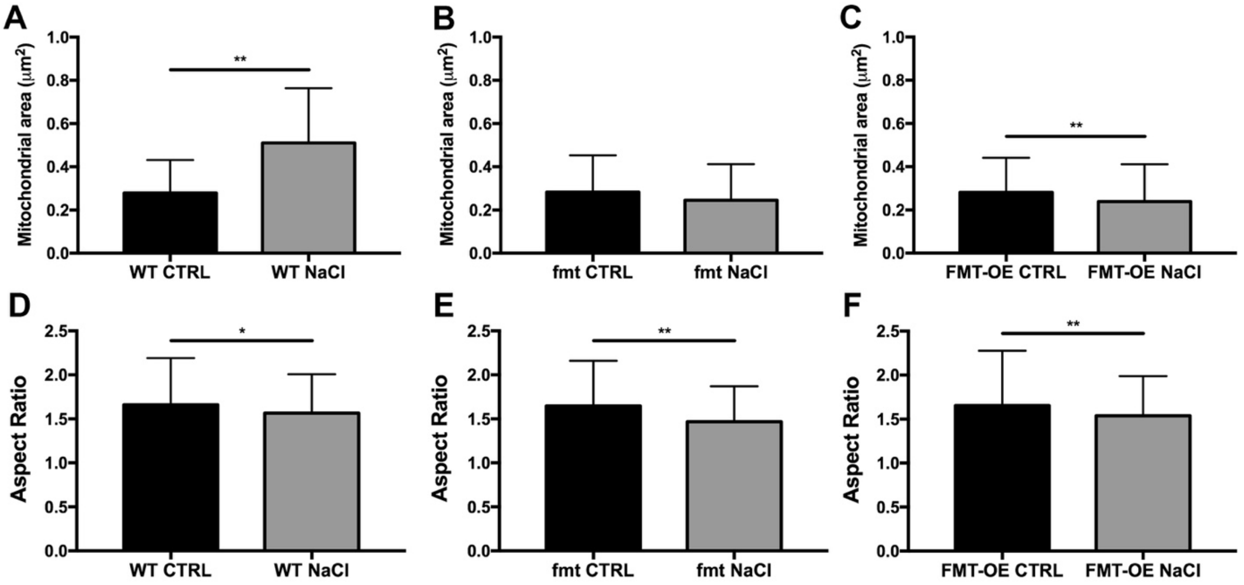

**Supplemental Figure S5**

**A**

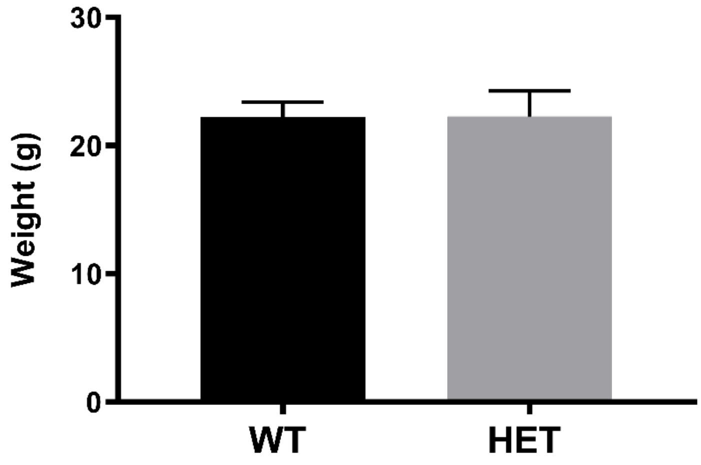


**B**

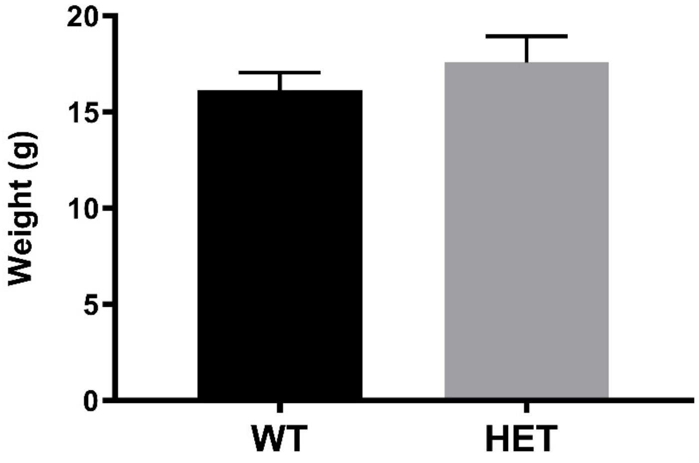
